# The G protein inhibitor YM-254890 is an allosteric glue

**DOI:** 10.1101/2024.11.25.625299

**Authors:** Tony Trent, Justin J. Miller, Gregory R Bowman

## Abstract

Given the prominence of G protein coupled receptors (GPCRs) as drug targets, targeting their immediate downstream effectors, G proteins, could be of immense therapeutic value. The discovery that the natural product YM-254890 (YM) can arrest uveal melanoma by specifically inhibiting constitutively active Gq/11without impacting other G protein families demonstrates the potential of this approach. However, efforts to find other G protein family-specific inhibitors have had limited success. Better understanding the mechanism of YM could facilitate efforts to develop other highly specific G protein inhibitors. We hypothesized that differences between the conformational distributions of various G proteins play an important role in determining he specificity of inhibitors like YM. To explore this hypothesis, we built Markov state models (MSMs) from molecular dynamics simulations of the Gα subunits of three different G proteins, as YM predominantly contacts Gα. We also modeled the heterotrimeric versions of these proteins where Gα is bound to the Gβγ heterodimer. We find that YM-sensitive Gα proteins have a higher probability of adopting YM-bound-like conformations than insensitive variants. There is also strong allosteric coupling between the YM- and Gβγ-binding interfaces of Gα. This allostery gives rise to positive cooperativity, wherein the presence of Gβγ enhances preorganization for YM binding. We predict that YM acts as an “allosteric” glue that allosterically stabilizes the complex between Gα and Gβγ despite the minimal contacts between YM and Gβγ.

## Introduction

The ability to target specific G protein families would be of immense therapeutic value. Heterotrimeric G proteins are intracellular switches that are activated by G protein-coupled receptors (GPCRs) to elicit biological responses by regulating downstream enzymes, ion channels or protein kinases^1–3^. Approximately one-third of currently approved drugs target GPCRs ^3^. Therefore, the discovery of inhibitors that directly target downstream proteins like G proteins can provide a novel approach to address limitations of traditional, GPCR-targeted drugs. Directly targeting G proteins is especially appealing in diseases caused by G protein mutations that result in constitutive activation, such as uveal melanoma and Sturge-Weber Syndrome, where GPCR inhibition would be ineffective^2,3^. Targeting the other G protein families (Gi, Gs, and G12/13) could also be useful in other settings^4,5^.

G proteins predominantly exist as multi-subunit complexes that are allosterically coupled in a manner that allows the components (especially the Gα subunit) to switch from inactive to active states in response to GPCR signaling (Figure 1) ^1–3^. In the inactive state, Gα subunits bind GDP and a Gβγ heterodimer. Activated GPCRs interact with surface elements of GDP-bound Gα subunits, including helix 5 (H5) and the loop between helix N and β-strand 1 (hNs1 loop)^6^. This interaction triggers allosteric conformational changes in Gα subunits up to 30 Å away from the GPCR-binding site, resulting in GDP and Gβγ release, GTP binding and initiation of downstream signaling^6,7^. Specifically, GPCR binding facilitates GDP release by causing disorder in the GDP-binding site, conformational changes in Switch I, and opening of the Ras-like and helical domains^7–9^. GPCR binding also causes conformational changes in Gα’s Gβγ-binding interface, notably Switch II, that promote dissociation of Gβγ from Gα. Subsequent binding of GTP stabilizes Gα subunits in their active state, which goes on to interact with other downstream components of the signaling cascade^9^.

**Figure 1.**
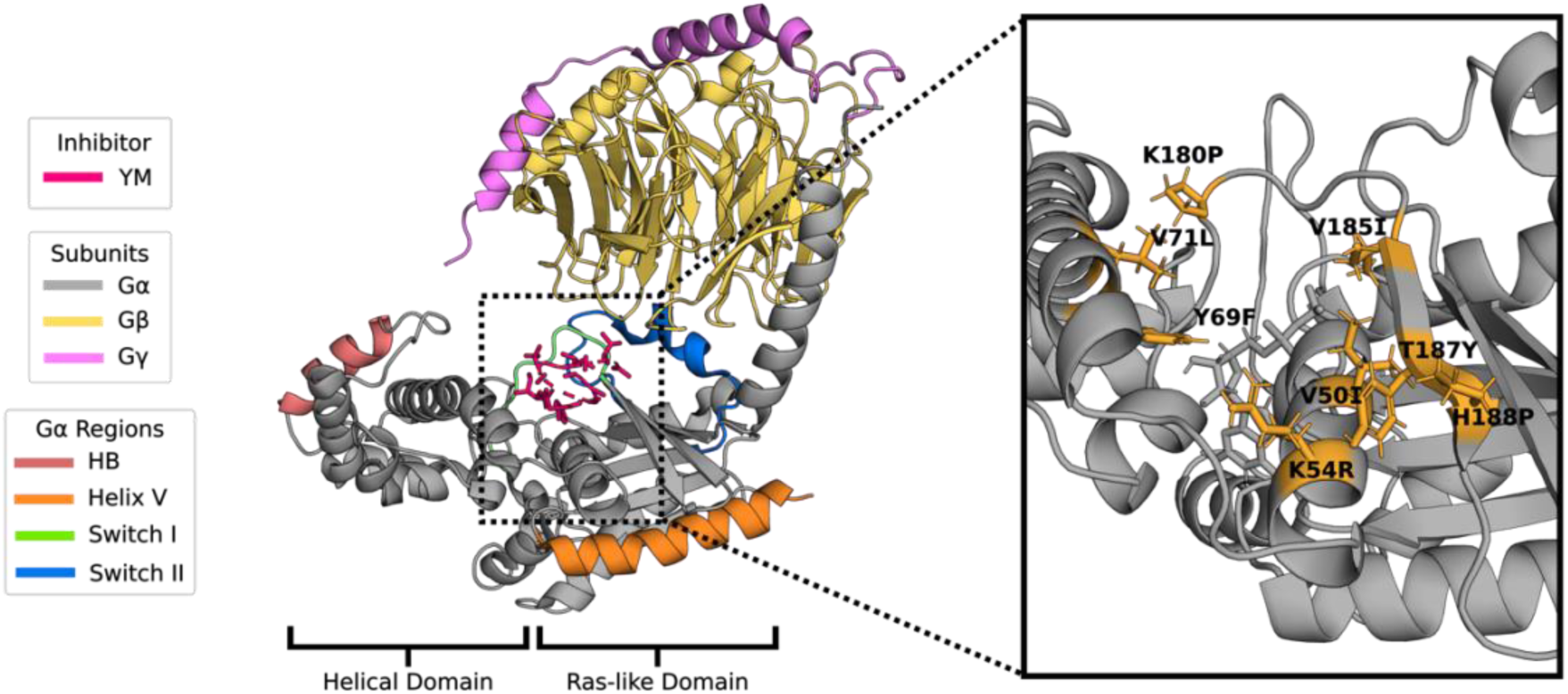
Structure of heterotrimeric Gq bound to the inhibitor YM-254890 (PDB ID: 3AH8)^6^. Relevant regions of the protein are colored according to the legend and residues that are mutated to create a YM-sensitive chimera of Gi1 are shown as sticks. The helical and Ras-like domains of the Gα subunit are also labeled.

A natural product called YM-254890^10^ (herein referred to as YM) is known to specifically inhibit the Gq/11 family of heterotrimeric G proteins but it has proved difficult to develop other compounds to specifically target Gq/11 or any of the other families without compromising potency^2,11,12^. YM is a macrocyclic, bacterial peptide that is thought to inhibit GDP release by blocking opening of the Ras-like and helical domains of Gα by binding at the hinge between these two domains (Figure 1)^8^. A closely related natural product from plants, called FR900359^13^ (FR), behaves similarly. To understand the mechanism of inhibition by YM, a YM-sensitive variant of Gi1 has been created by mutating 8 residues in the YM-binding site to match those in Gq^14^. This variant, called Giq, is more experimentally tractable than Gq and reproduces many properties of the Gq system, such as the greater sensitivity of the heterotrimeric form to YM than isolated Gα^14^. Using these resources to further understand the mechanism of YM inhibition could greatly accelerate efforts to develop more synthetically tractable inhibitors of Gq, as well as inhibitors of other G protein families.

Recent work has suggested that YM is a “molecular adhesive” (otherwise known as a glue) that stabilizes the GDP-bound off state by enhancing interactions between Gα and Gβγ^2^. This conclusion is primarily based on the observation that YM increases the melting temperature of the heterotrimer^15^. Other authors have also observed that YM is a far more potent inhibitor of heterotrimeric Gq than the isolated Gαq subunit^6,16^. YM’s IC50 for heterotrimeric Gq is somewherebetween 8 and 501nM^2^ but the IC50 for isolated Gα subunits is at least an order of magnitude weaker^6,14,16^. Interestingly, the authors note that YM only contacts one residue in Gβγ in available structures^6^. Furthermore, mutating this residue in Gβ only has a mild impact on YM’s potency^15^. Together, these data suggest there are additional mechanisms at work besides physical contacts.

Here, we explore the hypothesis that there is allosteric coupling between the binding of YM and Gβγ, such that YM acts as an allosteric glue whose ability to stabilize the G protein heterotrimer does not necessitate direct contacts with Gβγ. To explore this hypothesis, we used our simulation-based tools^11,17^ to assess whether there are differences between the structural preferences and allosteric networks of YM-sensitive and YM-insensitive G proteins.

Specifically, we ran all-atom simulations of Gq, Giq, and Gi1. We used these data to build maps of the conformational space of each G protein, called Markov state models (MSMs),^18^ and to infer the allosteric networks that allow each G protein to switch between active and inactive states. Then we compare the MSMs and allosteric networks for each variant to understand what makes some G proteins sensitive to YM and what makes others insensitive.

## Results

### Sensitive Gα isoforms have greater pre-organization for YM binding

Inspired by the conformational selection mechanism, we reasoned that YM-sensitive G protein isoforms may have a higher probability of adopting conformations that are well-suited to binding YM, which we call pre-organization. Experiments have demonstrated that both heterotrimeric Gq and the isolated Gαq subunit are sensitive to YM, while the heterotrimeric Gi1 and Gαi1 are not ^8^. We initially focus on the isolated α subunits as simulating them is less computationally demanding than simulating entire heterotrimers. In addition to simulating Gαq and Gαi1, we also chose to simulate Gαiq to see if this chimeric protein has more pre-organization than Gαi1.

To characterize the conformational ensembles of each Gα subunit, we ran all-atom molecular dynamics simulations with explicit solvent and constructed MSMs of each variant. Gαq simulations were started from the conformation of the α subunit taken from the full heterotrimeric YM-bound structure (PDB ID: 3AH8)^6^. Gαi1 and Gαiq simulations were started from homology models based on the starting structure used for Gαq. We ran 55-100 simulations initiated from each starting structure, for an aggregate of 98.55μs of simulation across the three Gα subunits. Then we built an MSM of each variant using Enspara^19^, with the conformational states determined by clustering the simulation data based on the Cα RMSD of residues surrounding the YM-binding pocket (specifically, residues within 5Å of YM in the YM-bound crystal structure^6^). We quantified pre-organization based on the distribution of RMSDs to the YM-bound crystal structure when weighted by the constructed MSMs.

As expected, we find that YM-sensitive isoforms have a reasonable probability of adopting conformations that are primed to engage YM while the insensitive isoform shows less pre-organization. The Cα RMSD distributions of the YM-binding pocket makes this distinction evident (Figure 2). Gαq and Gαiq are typically within 1-3 Å of the YM-bound structure. The similarity of them to the YM-bound structure even in the absence of the compound means that YM has to pay very little energetic cost to stabilize compatible protein structures over alternatives. In contrast, the insensitive Gαi1 typically adopts very different structures (3-5 Å from the YM-bound structure). That means there’s a large energetic cost for shifting probability density from these alternative structures to ones that are compatible with YM binding, which equates to a much weaker interaction with YM than more pre-organized variants have.

**Figure 2.**
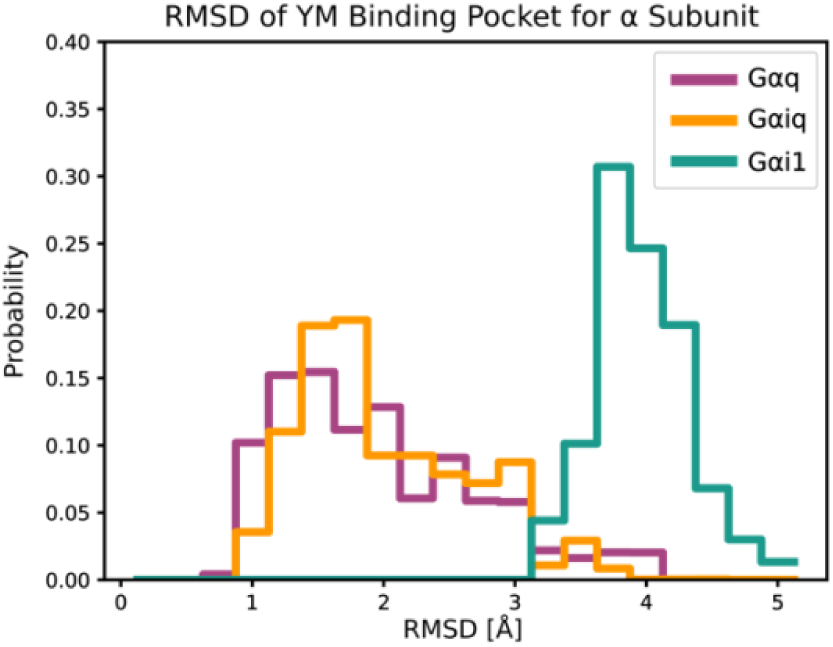
YM-sensitive Gα subunits have greater pre-organization for YM binding. Distributions of Cα RMSDs of the YM pocket in each isoform to a YM-bound crystal structure (PDB ID: 3AH8)^6^.

### Sensitive Gα isoforms display greater allosteric coupling between the YM-binding pocket and Gβ-binding interfaces

We hypothesized that the YM-binding pocket is allosterically coupled to the Gβγ-binding interface based on two experimental observations. First, YM is more potent of an inhibitor toward heterotrimeric Gq than Gαq^6,16^. Second, there are barely any contacts between Gβγ and YM in the available crystal structures^2,6^. The presence of allosteric coupling could provide a means for the presence of Gβγ to enhance Gαq’s affinity for YM, and vice versa, without there being significant direct contacts.

To explore this possibility, we sought to quantify the extent of allosteric coupling between the YM-binding pocket and several other regions of interest. In particular, we wanted to know if there is allosteric coupling between the YM-binding pocket and Switch II, as this key structural element is thought to undergo conformational changes during G protein activation and is the main binding interface for Gβγ. As reference points for judging the extent of this allosteric coupling, we also sought to quantify the allosteric coupling between the YM-binding pocket and two other regions known to be of importance for G protein activation, helix H5 and Switch I. As a negative control, we also sought to quantify the allosteric coupling between the YM-binding interface and helix HB. Our previous work found that this helix has little coupling to other parts of the protein and is not a key component of the allosteric network responsible for G protein activation^7^.

We used the correlation of all rotameric and dynamical states (CARDS) method to quantify the allosteric coupling between different regions of Gα and compare across isoforms^20^. CARDS uses the mutual information (MI) metric to quantify the information known about a given dihedral angle if you know the structure and dynamics of another. Then, the coupling between larger regions of a protein such as two different secondary structure elements can be obtained by summing the MI between all relevant pairs of dihedral angles. Comparing the total MI between two regions in two different protein variants provides a quick means to assess whether allostery is stronger between these regions in one variant in comparison to another.

As predicted, there is significant allosteric coupling between the YM-binding pocket and the Gβγ-binding interface in all three isoforms. (Figure 3A). Interestingly, Switch II in the YM-sensitive isoforms has significantly stronger coupling to the YM-binding interface when compared to the insensitive Gαi1. The greater pre-organization of the YM-binding pocket compounded with this observed stronger coupling could provide an explanation for the increase in YM affinity observed in the heterotrimeric forms of Gαq and Gαiq. The sensitive isoforms also have stronger coupling between the YM-binding pocket and the nearby Switch I region that is also thought to be important for G protein activation. This is expected as Switch I makes up the back of the YM binding pocket and makes significant contacts with YM when bound (Figure 1). However, the allosteric coupling of the YM-binding pocket to more distal regions of the protein is similar in all three proteins regardless of whether they are important components of the allosteric network that is responsible for G protein activation (e.g. helix H5) or not (e.g. helix HB).

**Figure 3.**
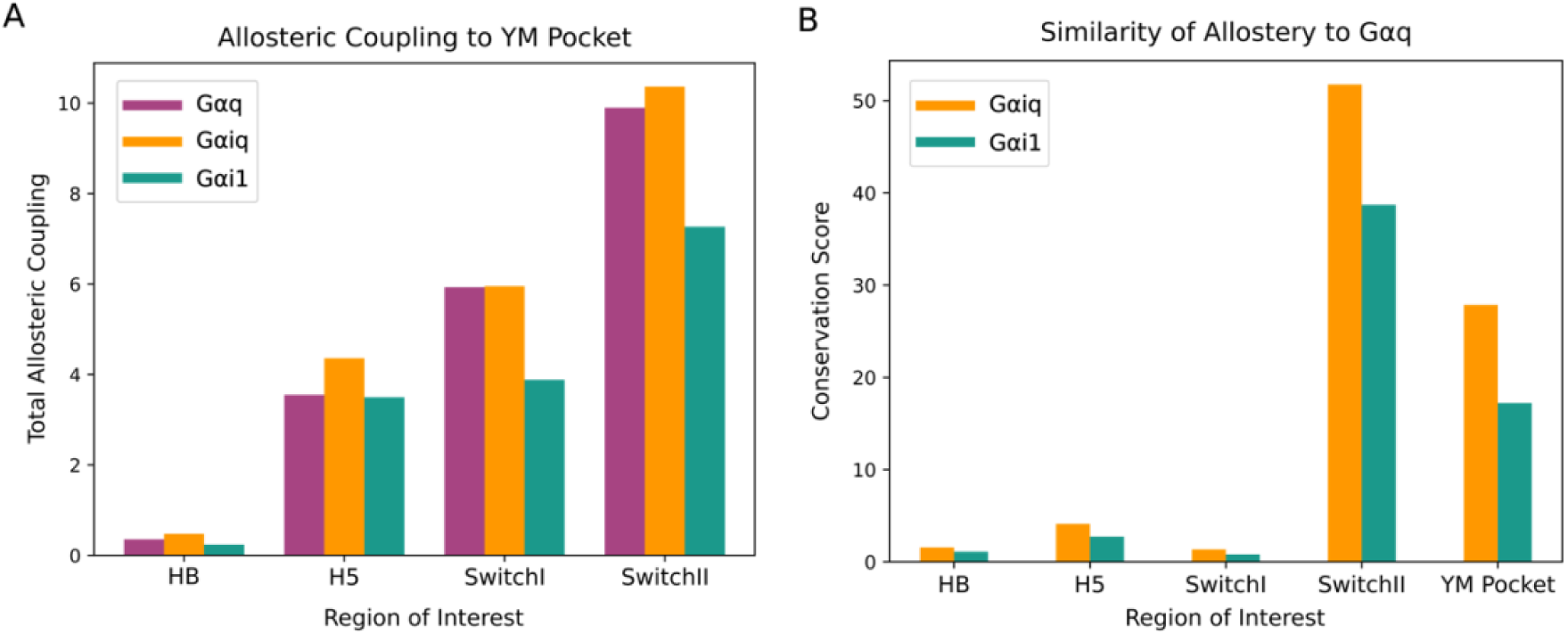
The YM-binding pockets of YM-sensitive Gα isoforms have stronger allosteric coupling to other key regions, notably switch II, which is a major part of the Gβγ-binding interface. A) The allosteric coupling between the YM-binding pocket and other parts of the protein. Switch II is of particular importance as it’s a major part of the Gβγ-binding interface. Switch I, Switch II, and H5 are all implicated in the allosteric network that is responsible for G protein activation. Meanwhile, HB is not strongly implicated in Gα’s allosteric network and is shown as a reference. Allostery is quantified using the CARDS method, as described in the Materials & Methods Section. B) Conservation of the allosteric networks observed in Gαiq and Gαi1 compared to Gαq. The conservation score reports on how similar allosteric coupling from a given region (e.g. H5) to all other parts of one isoform is to the coupling of that region in another isoform to all other parts of the protein.

The stronger allosteric coupling in Gαiq than Gαi1 between the YM-binding pocket and Switch II is a surprising find that warrants further investigation. The eight residues in Gαi1 that were mutated to match the sequence in Gαq were explicitly chosen because they directly interact with YM in the Gq crystal structure^6,14^. Since the residues mutated lack direct contact with Switch II, one might reasonably predict that the allosteric network in Gαiq would match that in Gαi1. Instead, we find that the extent of coupling between these two regions in Gαiq better matches that of Gαq. This begs the question of whether the Gαiq pocket mutations created some other allosteric connection that is absent in Gαi1or if Gαiq adopted an allosteric network more like that of Gαq.

To answer this question, we used an adapted network alignment method for assessing the total conservation of allostery between two proteins^21^. Instead of measuring the coupling between two specific regions like in Figure 2A, each isoform’s holistic allosteric network across the entire protein is compared to a corresponding reference network in another isoform. We chose to use Gαq as the reference to directly determine if Gαiq adopts a similar network. Then, like the coupling metric, we can then sum the conservation scores of residues into the same secondary structure elements to determine how the allosteric conservation of functional regions differs across isoforms.

The mutations in Gαiq remodel the protein’s allosteric network to more closely resemble Gαq than Gαi1 (Figure 3B). The allosteric conservation of the YM-binding interface and Switch II is well conserved in both Gαiq and Gαi1 when compared to the network of Gαq. However, the conservation is clearly heightened between Gαiq and Gαq. This trend extends for all regions tested on the protein including areas that showed no significant difference in coupling to the pocket (Figure 3A). The scores for other regions (HB, H5, and Switch I) are additionally much lower and the differences between Gαi1 and Gαiq compared to Gαq are much smaller. This difference suggests that there could be a selective pressure for the allosteric interaction between the YM-binding pocket and Switch II, which could be interesting to study in the future.

Together, our results strongly suggest positive cooperativity between YM and Gβγ binding to Gα. Fascinatingly, they also suggest that mutating the YM-binding pocket of Gαi1 to create the YM-sensitive Gαiq variant enhances allosteric coupling to the Gβγ-binding interface in a Gαq-like manner in addition to changing the direct interactions between the protein and YM^8^.

Previous reports suggest YM is a “molecular adhesive” (also known as a glue) that stabilizes interactions between the α subunit and the β subunit^2^. Our data suggests that the stabilization stems not only from the limited physical interactions between YM and the β subunit, but a pre-organization mechanism triggered allosterically from the pocket, expanding YM’s role to what we call an “allosteric glue”.

### Gβγ binding enhances pre-organization for YM binding in sensitive isoforms

To test whether the allosteric coupling we observed in the isolated Gα subunits enhances pre-organization in YM-sensitive isoforms, we also built models of the heterotrimeric forms of each G protein. We built starting structures for the sensitive isoforms based on the heterotrimeric crystal structure 3ah8 and the crystal structure 1gp2 for the insensitive Gi1^6,22^. We then ran approximately 31.86μs of aggregate simulation time across the three isoforms, constructed MSMs, and characterized the distribution of Cα RMSDs to the YM-bound structure of Gq. Including sidechains in the RMSD calculation is non-trivial because of how many sequence differences there are between the different protein variants. To account for differences in sidechain degrees of freedom, we compare the volume of YM-binding pocket between different isoforms. Comparing the distribution of pocket volumes for each isoform to the pocket volume in the YM-bound crystal structure provides a simple means to assess what fraction of the time each variant adopts conformations that can accommodate YM.

As predicted, Gβγ binding enhances pre-organization in the YM-sensitive heterotrimeric G proteins (Figure 4). As for the YM-binding pocket, Gq has the lowest RMSDs to the YM-bound crystal structure and Gi1 has the highest. However, the differences between the isoforms are much smaller. To further assess whether these differences are significant or not, we also examined the distribution of pocket volumes for each variant as a means to account for sidechain degrees of freedom. These pocket volume distributions show clear differences between the heterotrimeric proteins (Figure 4B). Heterotrimeric Gq is open enough to accommodate YM more than five times more often than heterotrimeric Gi1 and Giq falls in between.

**Figure 4.**
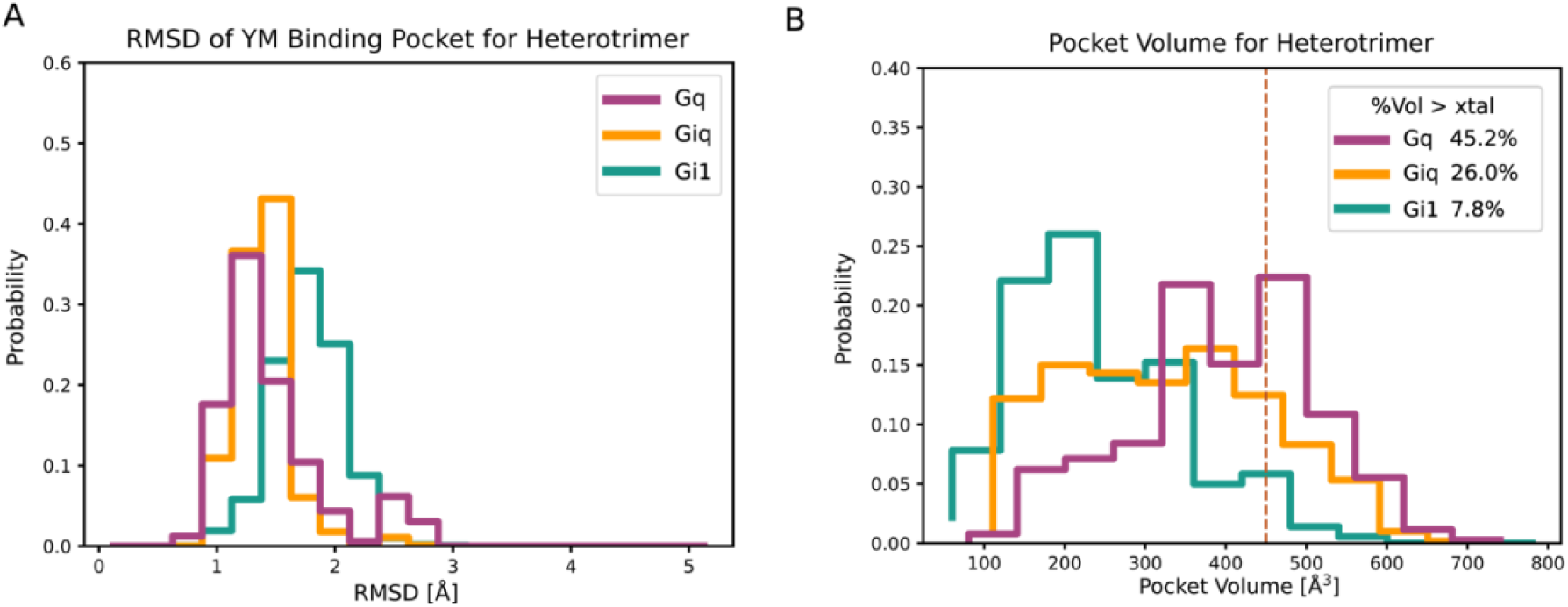
YM-sensitive heterotrimeric G proteins have greater pre-organization for YM binding. A) Distributions of Cα RMSDs of the YM pocket in each isoform to a YM-bound crystal structure ((PDB ID: 3AH8)^6^. B) Distributions of YM pocket volumes for each isoform. The pocket volume in the YM-bound crystal structure is shown as a vertical dashed line. The legend lists the percentage of the distributions that is greater than the volume in the crystal structure.

### Sensitive isoforms display more favorable binding to YM in both Gα and heterotrimeric forms

To test the impact of the pre-organization we have described on YM-binding affinity, we used the PopShift method to predict the dissociation constant (Kd) of YM for each isoform^23^. PopShift is a method for predicting binding free energies in a manner that accounts for protein conformational heterogeneity. Briefly, we estimated the Kd of YM for a representative structure for each state of an MSM using the GNINA software^24^. Then we used the probabilities of each state from our ligand-free MSM and the predicted Kd of YM for each state to estimate the net binding affinity of YM for the protein conformational ensemble. Importantly, the Kd values we predict with PopShift account for both the conformational preferences of the protein and the chemical interactions between YM and the protein. However, we note that docking algorithms are quite limited in their ability to quantitatively predict Kd values. We expect this approach can predict the rank order of YM’s affinity for different proteins but not YM’s absolute binding affinity for any of them.

As expected, we find that YM-sensitive isoforms have higher affinities (lower Kds) for YM (Figure 5). Both heterotrimeric Gi1 and the isolated Gαi1 subunits have much higher Kds for YM than the corresponding heterotrimeric or monomeric versions of Gαq and Gαiq. The predicted Kd of YM for heterotrimeric Gq is 27.76 nM, which is in reasonable agreement with the experimentally measured value of 10.96 nM [2]. However, the aforementioned limitations of molecular docking impacted the results of Gi1. The predicted Kd of YM for heterotrimeric Gi1 is 42.87 nM but experiments don’t show significant YM-binding even at uM concentrations of YM. While not quantitatively predictive, the method correctly captures that YM binds more strongly to sensitive variants than insensitive ones.

**Figure 5.**
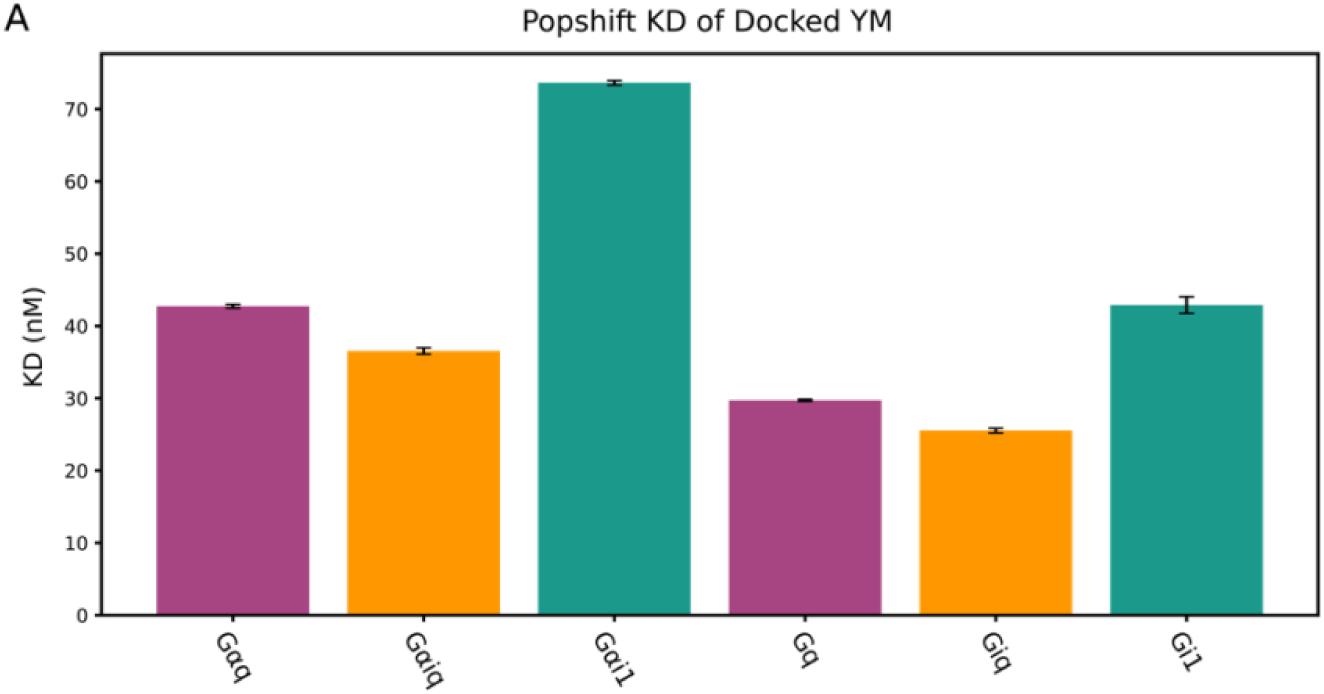
Sensitive isoforms have higher affinity for YM. Kd values were predicted using PopShift and error bars come from bootstrapping. The first three Kd’s are for isolated α subunits, while the last three are for heterotrimeric forms of each variant.

## Conclusions

G protein families perform a wide variety of functions in the cell and share considerable sequence and structural similarity, so targeting a specific family remains a significant challenge. A macrocyclic, bacterial peptide known as YM-254890 (YM) specifically targets and potently inhibits the Gq/11 family. However, YM’s mechanism of inhibition and the structural differences in G proteins that cause sensitivity to YM are largely unknown. We address this problem by using computational techniques to assess conformational behavior and relate that to specificity.

Here, we show that allostery has an important role to play in determining YM’s affinity and specificity, allowing it to act as an allosteric glue that stabilizes certain heterotrimeric G proteins without making significant contacts with all the subunits. Specifically, we found that the α subunits of sensitive isoforms have a preorganization mechanism where a binding competent YM-binding pocket is adopted preferentially (Figure 2). Additionally, we found that YM-sensitive isoforms have strong allosteric coupling between the YM- and Gβγ-binding sites whereas this coupling is weaker in insensitive isoforms(Figure 3). As a result of this allostery, the heterotrimeric versions of the sensitive isoforms have even greater pre-organization than their corresponding Gα subunits in isolation or any of the insensitive variants (Figure 4,5).

In the future, we expect that our tools and insights will be useful for informing the development of new G protein specific inhibitors.

## Methods

### Molecular Dynamics Simulations

Molecular dynamics simulations were seeded using x-ray crystallography structures (PDB IDs: 3ah8, 1gp2)^6,22^ and homology modelled mutants (Giq) for starting structures. The GROMACS^25^ software (version 2022.4) and the AMBER03 force field^26^ were used to carry out simulation. The proteins were each solvated using CHARMM-GUI^27,28^ in a dodecahedron box using TIP3P^29^ waters and 0.1 uM NaCl. Energy minimization was performed using steepest descent minimization, followed by NVT equilibration for 200 ps, NPT equilibration for 1 ns. Simulations were then either performed on the Folding@Home platform or locally^30^. Simulation times ranged from 100ns-500ns for an aggregate time of 98.55μs across the α-alone proteins and 31.86μs across the heterotrimeric proteins.

### Markov State Model (MSM) Construction

We clustered G protein conformations in trajectories and then used the resulting assignments to construct the Markov state model. First, we used Enspara^19^ and the hybrid k-center/k-medoids method algorithm on YM-pocket Cα atoms with a cutoff of 1.2Å. Pocket residues were determined by superimposing the YM bound crystal structure (PDB ID: 3AH8)^6^ onto the structure of interest and taking all residues within 5Å of YM. MSMs were then constructed using Enspara’s MSM builder^19^ using a lag time of 250ns. We then used bootstrapping for other analysis methods by reconstructing MSMs with trajectories dropped.

### LIGSITE

We used Enspara’s implementation of the LIGSITE algorithm to detect and quantify pocket volumes on the heterotrimeric G proteins^19,31^. Firstly, a cartesian grid was generated over the structure of interest. Then after determining which grid points are in pockets, grid points within a distance cutoff of the pocket residues are collected and converted into volumes. We used the hyperparameters of a probe radius of 1.4Å, a min rank of 5, a grid spacing of 0.7Å, a minimum cluster size of 3, and a distance cutoff of 5Å as previously described through our Enspara implementation of LIGSITE^19^. The only nonstandard choice of parameters was the min rank of 5 as previous uses of 6 were to find deep, cryptic pockets while the YM binding pocket is a more surface level, non-cryptic pocket.

### RMSD

RMSD calculations were performed on cluster centers using the MDTraj^32^ RMSD function and then weighted using the relative probability of each center as prescribed by the MSM.

### Correlation of All Rotameric and Dynamical States (CARDS) Analysis

To quantify allostery, we calculated mutual information between rotameric states of dihedral angles using a method called Correlation of All Rotameric and Dynamical States (CARDS)^20^. Categorizing dihedrals angles from both the side chain and backbone into greater rotameric states allows us to track an angle’s motions across the length of the simulation. We then used Mutual Information (MI) to assess the level of correlated motions between these metrics using the formula:

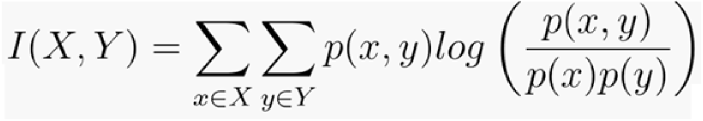

Where x ∈ X refers to all the possible states (x) that a dihedral (X) can adopt, p(x) is the probability that the dihedral adopts the specific state x, with the same holding true for state y and dihedral Y. Thus, p(x,y) is the joint probability that dihedral X adopts state x and dihedral Y adopts state y. This MI is then normalized using the formula:

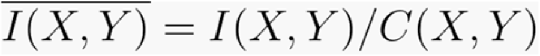

Where C(X,Y) represents the maximum possible MI between two dihedrals, called the channel capacity. This accounts for the possibility of certain angles having more possible states such as a sidechain versus a backbone.

To account for disordered communication, we additionally assigned states into ordered or disordered states based on whether they are stably in a rotameric state or transitioning between states respectively using the formula:

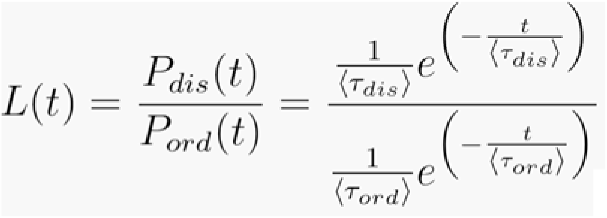

This formula is derivative from interpretations of Bayes factors, where P_dis_ is the probability of a state being disordered, P_ord_ is the probability of a state being ordered, and their respective tau values represent mean ordered times. State determination is disordered if L > 3.0 and ordered otherwise.

To capture the holistic correlation (I_H_), we can now use our normalized MI as such:

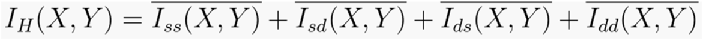

Where I_ss_ refers to the structure-structure correlation, I_sd_ refers to the structure-disorder correlation, I_ds_ refers to the disorder-structure correlation, and I_dd_ refers to the disorder-disorder correlation. The use of normalized MI allows for comparison between different types of correlations that can have different numbers of states. An example of this side-chain dihedrals, which have three possible rotameric states, so the maximum structural correlation (I_ss_) is log(3), but only two possible dynamical states so the maximum disordered correlation (I_dd_) is log(2).

### Molecular Docking

We used GNINA ^24^ to dock YM to the pockets of the G isoforms to determine a Kd for each pose. GNINA not only docks the ligand, but also uses a neural network to estimate Kds in a way that reflects experimental values more accurately. We then used Popshift to account for conformational heterogeneity of the proteins^23^. Popshift is a framework used to garner more accurate binding metrics produced by standard docking procedures using MSM reweighting. By using these techniques in tandem, we get probability weighted Kd values for poses obtained using the previously described clustering methods.

## Acknowledgements

We are grateful to the Folding@home community for helping generate simulation data. This work was supported by NIH NIGMS R35GM152085 (G.R.B.) and NSF MCB2218156(G.R.B.). We are grateful to Neha Vithani and Sukrit Singh for helping facilitate simulations and develop techniques used in our analyses.

## Competing Interests

The authors declare no competing interests.

## Notes

### Competing Interest Statement

The authors have declared no competing interest.

